# The Automated Systematic Search Deduplicator (ASySD): a rapid, open-source, interoperable tool to remove duplicate citations in biomedical systematic reviews

**DOI:** 10.1101/2021.05.04.442412

**Authors:** Kaitlyn Hair, Zsanett Bahor, Malcolm Macleod, Jing Liao, Emily S. Sena

## Abstract

**Background:** Researchers who perform systematic searches across multiple databases often identify duplicate publications. Identifying such duplicates (“deduplication”) can be extremely time-consuming, but failure to remove these citations can, in the worst instance, lead to the wrongful inclusion of duplicate data. Many existing tools are not sensitive enough, lack interoperability with other tools, are not freely accessible, or are difficult to use without programming knowledge. Here, we report the performance of our Automated Systematic Search Deduplicator (ASySD), a novel tool to perform automated deduplication of systematic searches for biomedical reviews.

**Methods:** We evaluated ASySD’s performance on 5 unseen biomedical systematic search datasets of various sizes (1,845 – 79,880 citations), which had been deduplicated by human reviewers. We compared the performance of ASySD with Endnote’s automated deduplication option and with the Systematic Review Accelerator Deduplication Module (SRA-DM).

**Results:** ASySD identified more duplicates than either SRA-DM or Endnote, with a sensitivity in different datasets of 0.95 to 0.99. The false-positive rate was comparable to human performance, with a specificity of 0.94-0.99. The tool took less than 1 hour to deduplicate all datasets.

**Conclusions:** For duplicate removal in biomedical systematic reviews, ASySD is a highly sensitive, reliable, and time-saving tool. It is open source and freely available online as both an R package and a user-friendly web application.

## Background

### What are duplicate publications?

Researchers performing a systematic review typically search across multiple biomedical databases to collect as many relevant citations as possible (1). This process can introduce a substantial number of duplicate citations (2). For example, overlap between EMBASE and PubMed is estimated to be as much as 79% (3). To further complicate matters, although publication of the same article in more than one journal is widely considered to be unethical (at least under most circumstances), we have identified many examples of this. In fact, six different patterns of duplicate publication have been identified (4) ranging from a direct “copy” of an article to so-called “salami” publications which slice up the data from one dataset into many resulting publications in an inappropriate manner (5). Such practices threaten scientific integrity and inflate redundancy in the literature (6). Different approaches may be required to tackle the distinct forms of “duplicate publication”.

Effective duplicate removal is an essential, if underappreciated, part of the data collection process of systematic reviews (7). If duplicate citations are not removed effectively, reviewers can waste time screening the same citations for inclusion, and run the risk of accidentally including the same paper more than once in their meta-analyses, leading to inaccurate conclusions (8). False positives (incorrect removal of citations which are not duplicates) can be just as problematic (9, 10) and reduce the accuracy and reproducibility of systematic reviews.

Here we consider the challenge of bibliographic duplicate detection – where the same publication in the same journal is retrieved from several biomedical databases. Current and common approaches to deduplication for systematic reviews are summarised in Table 1.

**Table 1:**
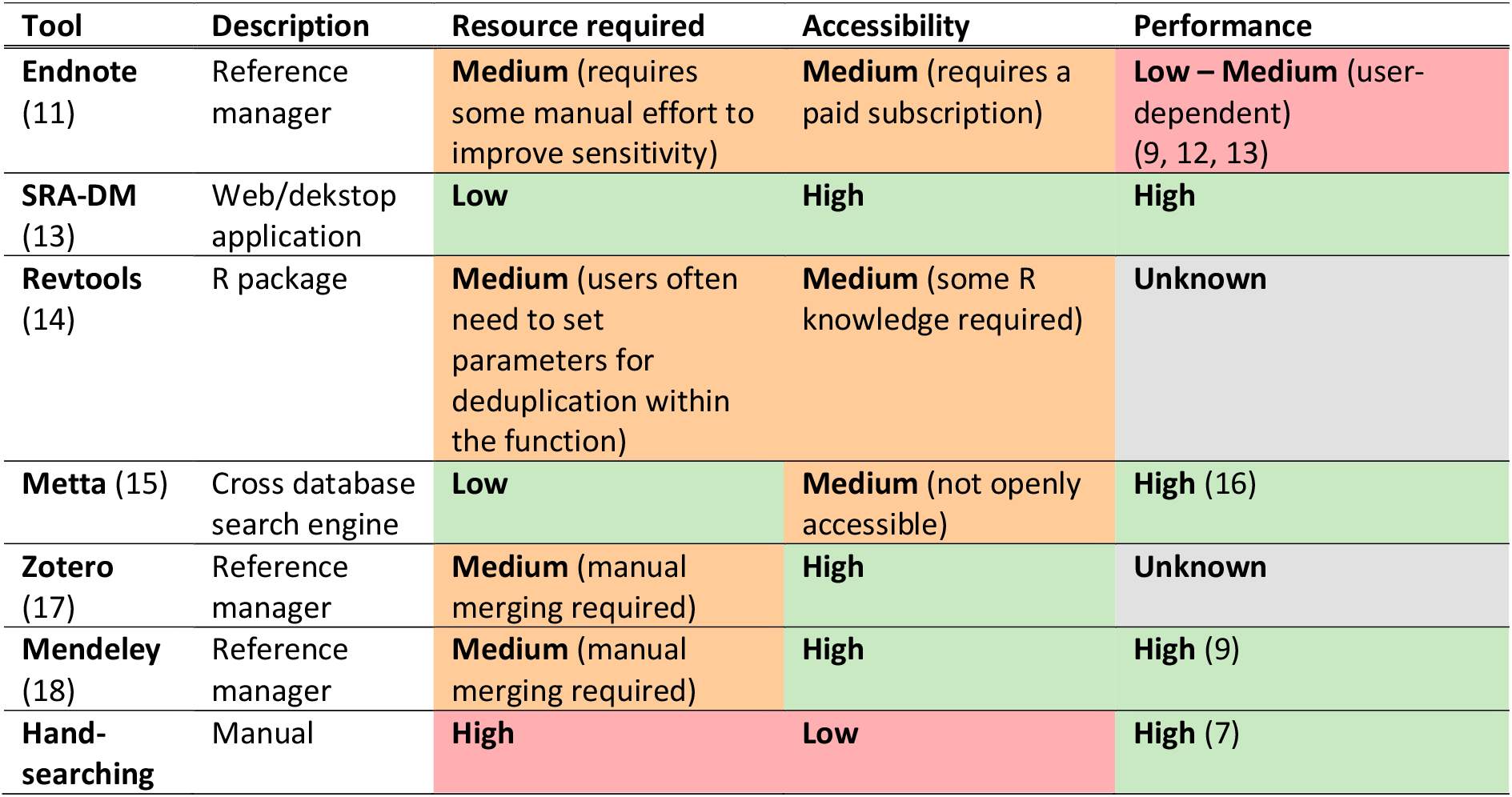
Current deduplication tools and approaches for systematic reviews.

| Tool | Description | Resource required | Accessibility | Performance |
| --- | --- | --- | --- | --- |
| **Endnote**<br>(11) | Reference manager | **Medium** (requires some manual effort to improve sensitivity) | **Medium** (requires a paid subscription) | **Low – Medium** (user-dependent)<br>(9, 12, 13) |
| **SRA-DM**<br>(13) | Web/desktop application | **Low** | **High** | **High** |
| **Revtools**<br>(14) | R package | **Medium** (users often need to set parameters for deduplication within the function) | **Medium** (some R knowledge required) | **Unknown** |
| **Metta** (15) | Cross database search engine | **Low** | **Medium** (not openly accessible) | **High** (16) |
| **Zotero**<br>(17) | Reference manager | **Medium** (manual merging required) | **High** | **Unknown** |
| **Mendeley**<br>(18) | Reference manager | **Medium** (manual merging required) | **High** | **High** (9) |
| **Hand-searching** | Manual | **High** | **Low** | **High** (7) |

### Identifying duplicates citations in preclinical systematic reviews

Systematic review findings often inform clinical practice. In recent years, largely in response to discrepancies between findings in laboratory research and clincial trial results, researchers have began to apply systematic review methodologies to summarise preclinical evidence from animal and cell models of disease (19, 20). When considering different tools to identify duplicates from preclinical systematic search datasets, we must take into account what type of citation data the tool was designed to deduplicate. For example, the databases supported by Metta are highly specific to clincial research and do not support search engines routinely used for preclinical reviews such as Web of Science. Furthermore, the type and extent of duplicate publications may differ in the preclinical literature – an author may publish a higher number of similar papers in a short space of time, or there may be less bibliometric information available for studies published in lesser known (and less frequently indexed) journals. Our group frequently retrieves tens of thousands of potentially relevant citations for a preclinical systematic review. Tools should therefore be evaluated on comparatively large datasets to determine the magnitude of gains and losses on that scale (e.g. how many duplicate citations a tool is likely to remove correctly). Previous evaluations of duplicate removal tools have used relatively small (<5,000 citations) systematic search datasets primarily representing clinical research citations (2, 9, 13).

### Current methods of duplicate removal

Researchers often use citation managers to remove duplicates, as these are easy to use and straightforward to integrate into typical systematic search methods. Among them, Endnote is one of the most established (21). Endnote’s “find duplicates” feature automatically detects citations matching on Author, Year and Title by default. Users can also adjust the match criteria within Endnote’s settings (i.e. match on Title and Journal) to identify additional duplicate records. The requirement for a 100% match to identify duplicates, however, results in many records being missed. Small differences in the way the Titles, Authors, and Journals are represented are extremely common. Deduplication might be simplified through the use of unique identifiers for journal articles such as PubMed IDs (PMIDs) or digital object identifiers (DOIs). However, Endnote does not provide an option within their deduplication settings to match citations based on DOIs, PMIDs, Accession Numbers, or URLs. Matching is further complicated by indexing differences in the formatting of page ranges, with some biomedical databases adopting a longer form (1234-1235) and some a shorter form (1234-5); although an import filter has been developed to address this issue in Endnote (22).

Endnote’s auto-deduplication feature is an attractive option due to its simplicity, yet there is a wealth of evidence to suggest it is an imperfect solution; as it fails to identify more duplicates (higher number of false negatives) and removes more citations incorrectly (higher number of false positives) than other citation managers (9). Moreover, our prior experience of using Endnote is that many duplicates remain in large datasets even after extensive deduplication using a combination of automated and user-configured methods.

Many citation managers, including Endnote, are proprietary software which restricts their accessibility, prevents intereroperability, and limits transparency about how their underlying duplicate detection process works. Increasingly, freely available open-source citation managers such as Zotero and Mendeley have gained popularity. Both have integrated deduplication tools which match citations automatically, then require users to manually select citations to merge within each matching group.

Several other tools for duplicate removal have emerged in recent years, either as stand-alone tools or as part of alternative workflows (which may bypass the need for traditional citation managers). The “Systematic Review Assistant” (SRA) is a suite of free, open-source systematic review tools developed by researchers at Bond University. Their “deduplication module” (SRA-DM) has a user-friendly interface in which users can upload a search file in various formats and perform automated duplicate removal in a few clicks. SRA-DM has been shown to identify substantially more duplicates than Endnote (13, 16). Another option is the metasearch engine Metta, which automatically removes duplicate citations appearing across 5 medical databases including PubMed, EMBASE, CINAHL, PsycINFO and Cochrane Central Register. One can also de-duplicate using Revtools, an R package. Of course, manual deduplication is strongly advised to complement these automated approaches (2), but this is time consuming and can lead to errors (9).

### Deduplication tools to support “Living” or automated reviews

Increasingly, meta-researchers are aspiring to provide automated or “living” systematic reviews (23), producing real-time summaries of a domain including the most recent research findings. To enable such summaries, we need automation tools at each stage that are reliable and require minimal manual intervention. Where review teams are large, as is the case in crowdsourced reviews, the risk of duplicate studies being retained is likely higher. Sensitivity of a deduplication tool (ability to detect duplicates) is therefore of paramount importance, since several reviewers could extract information from a given paper, unaware that others were also doing so. Furthermore, if machine learning approaches are used to select included studies, duplicate publications present in the training data may reduce the performance of classifiers.

Deduplication tools should be interoperable and easily integrated into automated workflows. Tools with a programmatic component are likely superior in this respect because once they have been configured, they may be implemented in a data pipeline without manual intervention. Depending on the project goals, it may be useful to have some control over the tool’s duplicate removal logic. For instance, if two records are identified to be duplicates of each other, which record should be retained? It may be useful to configure the tool to retain the existing version of a citation when a new, matching citation is identified in an updated search, so that existing annotations and data extractions can be retained. This approach could also be used in more conventional systematic review updates, often occurring after many years (24) and often involving significant overlap between systematic search dates to prevent missing relevant studies. Alternatively, researchers may wish to preferentially retain the newer citation, which may be more complete and may contain more accurate meta-data.

We developed the ASySD to identify and remove bibliographic duplicates from preclinical systematic review searches. The tool allows users to label which reviews should be preferentially maintained (e.g. older citations); and can be accessed either through a web application or integrated programmatically with automated workflows via an R package. We critically evaluated ASySD in comparison with two user-friendly, low effort automated tools - Endnote’s automated duplicate removal and Bond University’s SRA-DM.

## Methods

Prior to performance evaluation we registered a protocol describing our methods on the Open Science Framework (25).

### Definition of “Duplicate citations”

We define bibliographic duplicates as the presence of two or more citations representing the same publication within an aggregated systematic review search result, even where those citations differ subtly in recorded details such as author(s), title, journal pagination, issue number or volume. If the same study is published in two separate journals, we do not consider this a duplicate citation for these purposes. Similarly, sets of conference abstracts, preprints and journal articles which describe the same research are not be classed as duplicate citations.

### Tool development and functionality

We developed ASySD in the R programming language. To improve the chance of detecting duplicate citations, data undergoes several cleaning and formatting steps. This includes renaming missing or anonymous Authors as “Unknown”, harmonising differences in DOI format, removing punctuation, and making all citation information upper case.

Using the RecordLinkage R package (26), we applied blocking criteria (fields which must be a 100% match) to identify possible duplicate pairs. These criteria were largely based on guidance to systematically identify all possible duplicates using Endnote’s manual 100% match filters (22). Blocking criteria (see Table 2) were applied in four separate rounds because of the extensive memory requirements needed to perform these operations on large datasets in R; however, matches identified within any of the rounds were considered a possible duplicate pair.

**Table 2:** Blocking criteria specified for ASySD to identify potential duplicate citations.

| Order | Blocking criteria (100% match on specified fields) |
| --- | --- |
| Round 1 | (Title AND Pages) OR<br>(Title AND Author) OR<br>(Title AND Abstract) OR<br>DOI |
| Round 2 | (Author AND Year AND Pages) OR<br>(Journal AND Volume AND Pages) OR<br>(ISBN AND Volume AND Pages)<br>(Title AND ISBN) |
| Round 3 | (Year AND Pages AND Volume) OR<br>(Year AND Issue AND Volume) OR<br>(Year AND Pages AND Issue) |
| Round 4 | (Author AND Year) OR<br>(Title AND Year) OR<br>(Title AND Volume) OR<br>(Title AND Journal) |

Most pairs identified with blocking criteria are not true duplicates, and further comparisons are needed to ascertain duplicate status. To compare the overall similarity of a matching pair, we also calculate string comparisons across all relevant fields (Title, Year, Journal, ISBN, Abstract, DOI, Issue, Pages, and Volume) using the RecordLinkage package. Using a heuristic approach, we developed and applied additional match filters based on string comparison match strength (a numerical value between 0 and 1) to optimise performance and prevent the deletion of citations which were not duplicates. During development, we used three existing CAMARADES systematic review search results with labelled duplicates (Neuropathic Pain (27), Antioxidants (28), and Epilepsy (29)) to iteratively validate and adjust the match filters to improve the performance of the tool.

Once ASySD identifies all matching citations, one citation is removed from each pair. First, citations which do not contain abstracts are preferentially removed. Where a newer version of a citation exists (e.g. e-publication date versus publication date), we will preferentially retain the most up-to-date version. If neither of these rules apply (e.g. both citations contain abstract text, and have the same year of publication), then the second listed citation in each pair is removed. Where there are more than two duplicates, the code logic ensures that only one is kept from within each duplicate set. There is an option for users to set a preference for citations to be retained in the dataset using a “Label” field. If specified, duplicates are ordered so that these citations are always the first citation in each pair, and are therefore retained.

Citation pairs which fall short of the additional match filters but still have high string comparison scores are retained for manual deduplication – where users can manually review these matches and select which (if any) citation of the two they would like to remove from the search.

The underlying code for ASySD is open-source and available on Github, where it is also available to download as an R package (30). To ensure accessibility, we have also created a user-friendly web application build using R Shiny (31). Users can upload a file with search returns (e.g. Endnote .xml, .csv, or .txt file), click a button to run the deduplication procedure, complete any additional manual deduplication within the application (if required), and download the results as a .csv file or a tab delimited .txt file (formatted for importing into Endnote). For transparency, there is the option to download a file with the all potentially matching pairs side-by-side (from initial blocking criteria) and to download all matching pairs after the additional filters were applied. The code underlying the Shiny web application is also available on Github (32).

### Gold-standard systematic search datasets

We assessed the performance of automated deduplication tools on five test datasets of varying sizes from systematic review searches (Table 3). For each dataset, duplicate citations had been removed in Endnote using a combination of automated deduplication functions, changing field parameters to identify all citations which match on certain field e.g. “Title”, and manual checking. Citations which had been removed by the human reviewer were reinstated and labelled as duplicates. We obtained three systematic search datasets from external sources, described below. We also used two datasets curated as part of ongoing in-house projects, a systematic review of systematic reviews of animal models of human disease (SRSR), and a systematic review of animal models of depression.

**Table 3:** Gold standard systematic search datasets.

| Dataset description | Databases searched | Citations obtained | Duplicates removed | Citations remaining |
| --- | --- | --- | --- | --- |
| **Diabetes dataset:** Antidiabetics in animal models of atherosclerosis (SYRCLE, Radboud University) (33) | Pubmed, EMBASE | 1,845 | 896 | 949 |
| **Neuroimaging dataset:** Epigenetic neuroimaging (MRC Centre for Reproductive Health, University of Edinburgh) (34) (*Preclinical (in vivo) and clinical data included in review*) | SCOPUS, EMBASE, Medline, Web of Science, | 3,438 | 1,280 | 2,158 |
| **Cardiac dataset:** Efficacy of cardiac ischemic preconditioning in animal models (SYRCLE, Radboud University) (35) | Pubmed, EMBASE | 8,948 | 3,153 | 5,795 |
| **Depression dataset:** Preclinical animal models of Depression (CAMARADES, University of Edinburgh) (36) | PubMed, EMBASE, Web of Science, | 79,880 | 9,418 | 70,462 |
| **Systematic review of systematic reviews (SRSR) dataset:** Systematic review of preclinical systematic reviews dataset (CAMARADES, University of Edinburgh) (37) | PubMed, EMBASE, Web of Science, | 53,001 | 16778 | 36223 |

Importantly, none of these datasets had been used in the development of the tool. To assess the time taken to perform “gold-standard” deduplication, we measured the time taken to deduplicate the SRSR dataset. To identify duplicates, we imported the systematic search into Endnote and followed recommended guidance (22) to systematically identify all duplicate citations in the dataset using a range of different matching field parameters e.g. matching on “Author” and “Year”.

### Methods for performance evaluation in testing datasets

To obtain the most up-to-date citation information and ensure all systematic searches for validation have a similar depth of information, we used the “find reference updates” feature in Endnote X9 to retrieve additional information (e.g. DOIs, page numbers, issue numbers, journal volumes).

We compared the performance of the ASySD tool (automated, with no manual input, deduplication mode only), Endnote X9 automatic deduplication, and SRA-DM (13) on the five gold-standard search datasets. To assess auto-deduplication performance using Endnote X9, we auto-deduplicated citations based on “author”, “year” and “title” matching criteria and using the “ignore spacing and punctuation” feature. In SRA-DM, we uploaded XML files of our datasets to the offline version of the tool (as the server has limited capacity for high volume datasets) and chose the automated deduplication option to remove all suspected duplicates. In the ASySD tool, we uploaded citations as an XML file to the web application and ran automated deduplication. Because of memory limitations on the shinyapps.io server, for search results containing over 50,000 citations, we ran the R Shiny application locally in R

To preferentially retain records which had been labelled as duplicates by the human reviewer (so that we would know that these had been identified as duplicates), we used the “labelled duplicates” feature of ASySD to preferentially remove citations which the human had also removed. Importantly, this process does not affect the accuracy of the tool – only the choice of which citation from each pair is removed. This made the deduplication process of ASySD as similar as possible to support that of the human reviewer, and provide a fair test of the performance of the tool.

Once duplicates were removed using each of the other tools, a “Duplicate ID” was generated for matching sets of duplicates identified by ASySD. This was possible because ASySD allows users to download the Record IDs of matching citation pairs. For each Duplicate ID there should therefore be one single citation labelled as “KEEP” and the remainder (one or more duplicate citations) labelled as “REMOVE”. We carried out extensive manual checking in MS Excel to interrogate duplicate citations identified by some approaches but missed by others, to ensure that they were indeed duplicates. We manually searched to identify additional studies and corrected the Duplicate ID as appropriate. All data (including the original de-duplicated search datasets, results from each deduplication tool, final manually checked datasets with duplicate IDs, and the R code used to assess performance) are available to view on the Open Science Framework (DOI: 10.17605/OSF.IO/C9EVS). Once each search file had been corrected, we analysed this final dataset in R to calculate performance.

We report the performance of each tool by calculating:

- Number of true positives (citations which are duplicates which are correctly removed from the dataset);
- Number of false positives (citations which are not duplicates which are wrongly removed from the dataset);
- Number of true negatives (citations which are not duplicates which correctly remain in the dataset);
- Number of false negatives (citations which are duplicates which remain in the dataset but which should have been removed).
- 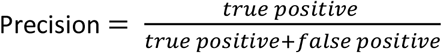
- 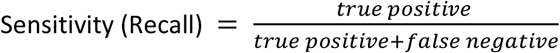
- 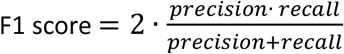

We also recorded any duplicates found by any of these approaches which had not been identified by humans in our “gold standard” datasets. We also recorded the time taken by each tool to deduplicate each dataset.

## Results

### True duplicates identified by any method

Across all datasets, additional duplicates were identified by automated tools which had been missed by the human reviewer(s). Furthermore, a small number of citations had been removed incorrectly by the human reviewer(s). We carefully considered all discrepancies between human reviewers and the automated tools to derive a new “gold standard” annotation against which to compare all approaches.

### Diabetes dataset

The Diabetes dataset (N=1,845) had 1,261 duplicate citations (68.3% of total; Table 4), of which 896 had been identified by human reviewers in the course of the systematic review, and a further 368 identified by at least one of the automated approaches and later confirmed by human scrutiny. While the sensitivity of the human approach was low, the specificity was high; only three citations were removed which were not duplicates (Table 5). Endnote, the SRA-DM, and ASySD were highly sensitive (sensitivity = 0.966, 0.910, and 0.998 respectively), but SRA-DM had a higher rate of false positives (n=70 citations incorrectly removed). The ASySD tool outperformed all other automated methods in terms of sensitivity (0.998), specificity (1.0), precision (1.0), and F1 score (0.999). Each automated deduplication method took less than 5 minutes to identify and remove duplicates in the diabetes dataset.

**Table 4:** Record classification in the Diabetes dataset by each deduplication method.

| Deduplication method | Duplicate citations removed | Citations remaining |
| --- | --- | --- |
| **TRUE duplicates**<br>(all methods + hand searching) | 1,261 | 584 |
| **Human** | 896 | 949 |
| **Endnote (automatic)** | 1,218 | 627 |
| **SRA-DM** | 1,217 | 628 |
| **ASySD** | 1,259 | 586 |

**Table 5:** Performance of deduplication tools in the Diabetes dataset.

|  | True + | True - | False - | False + | Sensitivity | Specificity | Precision | F1 | Time |
| --- | --- | --- | --- | --- | --- | --- | --- | --- | --- |
| **Human** | 893 | 581 | 368 | 3 | 0.708 | 0.995 | 0.997 | 0.828 | Unknown |
| **Endnote** | 1,218 | 584 | 43 | 0 | 0.966 | 1.0 | 1.0 | 0.983 | <5 minutes |
| **SRA-DM** | 1,147 | 514 | 114 | 70 | 0.910 | 0.880 | 0.942 | 0.926 | <5 minutes |
| **ASySD** | 1,259 | 584 | 2 | 0 | 0.998 | 1.0 | 1.0 | 0.999 | <5 minutes |

### Neuroimaging Dataset

The Neuroimaging dataset (N = 3,434) had 1293 duplicate citations (37.2% of total; Table 6). In this dataset, the human reviewer was highly sensitive and identified the vast majority of duplicate citations (sensitivity = 0.985; Table 7). However, a few citations had been removed in error (n=6), and a small number of duplicate citations were missed (n=19). Automated deduplication by Endnote and the SRA-DM was lacking in sensitivity and each missed hundreds of duplicates (n=310 and 243 respectively). The SRA-DM incorrectly removed a substantial number of citations (n=42). The false positives rate of the ASySD (n=4) and Endnote (n=3) were comparable to human performance.

**Table 6:** Record classification in the Neuroimaging dataset by each deduplication method.

| Deduplication method | Duplicate citations removed | Citations remaining |
| --- | --- | --- |
| **TRUE duplicates**<br>(all methods + hand searching) | 1,293 | 2,145 |
| **Human** | 1,280 | 2,158 |
| **Endnote (automatic)** | 986 | 2,452 |
| **SRA-DM** | 1,092 | 2,346 |
| **ASySD** | 1,282 | 2,156 |

**Table 7:** Performance of deduplication tools in the Neuroimaging dataset.

|  | True + | True - | False - | False + | Sensitivity | Specificity | Precision | F1 | Time |
| --- | --- | --- | --- | --- | --- | --- | --- | --- | --- |
| **Human** | 1,274 | 2,139 | 19 | 6 | 0.985 | 0.997 | 0.996 | 0.990 | Unknown |
| **Endnote** | 983 | 2,142 | 310 | 3 | 0.760 | 0.999 | 0.997 | 0.863 | <5 minutes |
| **SRA-DM** | 1,050 | 2,103 | 243 | 42 | 0.812 | 0.980 | 0.962 | 0.880 | <5 minutes |
| **ASySD** | 1,278 | 2,141 | 15 | 4 | 0.988 | 0.998 | 0.997 | 0.993 | <5 minutes |

Overall, the ASySD tool outperformed all other automated methods in terms of sensitivity (0.998), specificity (0.998), precision (0.997), and F1 score (0.993). Each method took under 5 minutes to identify and remove duplicates.

### Cardiac Dataset

This cardiac dataset (N = 8,948) contained 3,510 duplicate citations (39.2% of total; Table 8). The human reviewer sensitivity was high, and they captured most duplicates (sensitivity = 0.893; Table 9). Seventeen records had been removed in error. Endnote missed a substantial portion of duplicates (sensitivity = 0.749). The SRA-DM identified many false positives (n=275) and missed many duplicates (n=2,361). The ASySD tool outperformed other automated methods in terms of sensitivity (0.998) and F1 score (0.998) and was matched by Endnote in specificity (0.999) and precision (0.999). Deduplication took less than 5 minutes using Endnote or ASySD and just under 30 minutes using the SRA-DM.

**Table 8:** Record classification in the Cardiac dataset by each deduplication method.

| Deduplication method | Duplicate citations removed | Citations remaining |
| --- | --- | --- |
| **TRUE duplicates**<br>(all methods + hand searching) | 3,510 | 5,438 |
| **Human** | 3,153 | 5,795 |
| **Endnote (automatic)** | 2,737 | 6,211 |
| **SRA-DM** | 1,424 | 7,524 |
| **ASySD** | 3,507 | 5,441 |

**Table 9:**
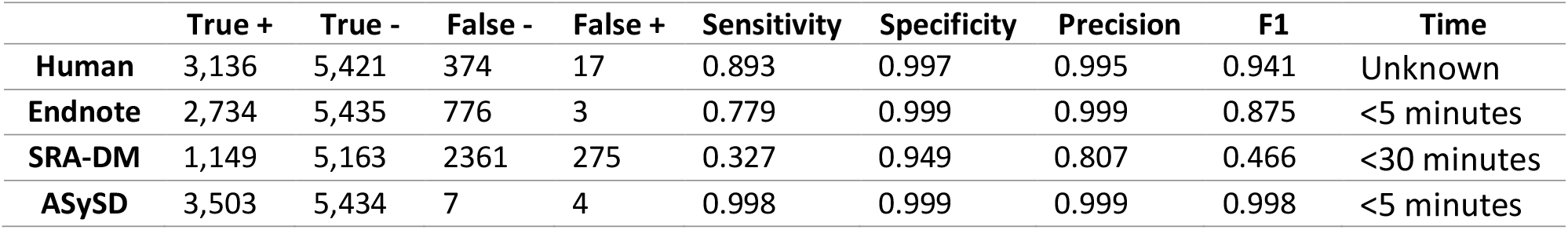
Performance of deduplication tools in the Cardiac dataset.

### Depression Dataset

The depression dataset (N=79,880) contained 10,059 duplicate citations (12.6% of total; Table 10). The human reviewer sensitivity was very high, and they correctly identified most duplicates. Endnote missed many duplicate citations (sensitivity = 0.75; Table 11) but was highly specific, removing only five duplicate citations incorrectly. The SRA-DM was highly sensitive (sensitivity = 0.98) but removed a substantial number of false positive duplicates (n=1,348). Overall, ASySD had a higher sensitivity (0.957), specificity (0.999), precision (0.993) and F1 score (0.974) than other automated tools. Deduplication using Endnote or ASySD took less than an hour, while the SRA-DM took approximately 48 hours to complete the process.

**Table 10:** Record classification in the Depression dataset by each deduplication method.

| Deduplication method | Duplicate citations removed | Citations remaining |
| --- | --- | --- |
| **TRUE duplicates**<br>(all methods + hand searching) | 10,059 | 69,821 |
| **Human** | 94,18 | 70,462 |
| **Endnote (automatic)** | 75,36 | 72,344 |
| **SRA-DM** | 10,796 | 69,084 |
| **ASySD** | 96,96 | 70,184 |

**Table 11.** Performance of deduplication tools in the Depression dataset.

|  | True + | True - | False - | False + | Sensitivity | Specificity | Precision | F1 | Time |
| --- | --- | --- | --- | --- | --- | --- | --- | --- | --- |
| **Human** | 9,390 | 6,9793 | 669 | 28 | 0.933 | 0.999 | 0.997 | 0.964 | Unknown |
| **Endnote** | 7,531 | 6,9816 | 2,528 | 5 | 0.749 | 0.999 | 0.999 | 0.856 | <30 minutes |
| **SRA-DM** | 9,448 | 6,8473 | 611 | 1,348 | 0.939 | 0.980 | 0.875 | 0.906 | ~48 hours |
| **ASySD** | 9,624 | 6,9749 | 435 | 72 | 0.957 | 0.999 | 0.993 | 0.974 | <1 hour |

### Systematic review of systematic reviews dataset

The SRSR dataset (N=53,001) had 16,838 duplicate citations (31.7% of total; Table 12). The human reviewer sensitivity was high (sensitivity = 0.990; Table 13), capturing nearly all duplicates and outperforming other methods. Endnote lacked sensitivity (0.760) and removed the fewest citations overall. The SRA-DM identified many false positives (n=1868) and lacked sensitivity (0.709). The ASySD tool outperformed other automated methods in terms of sensitivity (0.982), precision (0.999) and F1 score (0.991) and was matched by Endnote on specificity (0.999), with a low false positive rate. Manual deduplication had taken one team member (ZB) approximately 9 hours to complete using Endnote. Automated deduplication via ASySD and Endnote took less than 1 hour, and the SRA-DM took just under 24 hours.

**Table 12:** Record classification in the SRSR dataset by each deduplication method.

| Deduplication method | Duplicate citations removed | Citations remaining |
| --- | --- | --- |
| **TRUE duplicates**<br>(all methods + hand searching) | 16,838 | 36,163 |
| **Human** | 16,778 | 36,223 |
| **Endnote (automatic)** | 12,830 | 40,171 |
| **SRA-DM** | 13,814 | 39,187 |
| **ASySD** | 16,564 | 36,437 |

**Table 13:** Performance of deduplication tools in the SRSR dataset.

|  | True + | True - | False - | False + | Sensitivity | Specificity | Precision | F1 | Time |
| --- | --- | --- | --- | --- | --- | --- | --- | --- | --- |
| **Human** | 16,668 | 36,053 | 170 | 110 | 0.990 | 0.997 | 0.993 | 0.992 | ~9 hours |
| **Endnote** | 12,794 | 36,127 | 4,044 | 36 | 0.760 | 0.999 | 0.997 | 0.862 | <1 hour |
| **SRA-DM** | 11,946 | 34,295 | 4,892 | 1,868 | 0.709 | 0.948 | 0.865 | 0.779 | <24 hours |
| **ASySD** | 16,543 | 36,142 | 295 | 21 | 0.982 | 0.999 | 0.999 | 0.991 | <1 hour |

### Overall performance

Across all datasets, Endnote’s automated deduplication function and ASySD had consistently low false-positive rates and high specificity. ASySD correctly identified more duplicate citations than Endnote (and often more than the human reviewer). SRA-DM removed more duplicates than Endnote in some cases, but the false-positive rate of SRA-DM was high. Compared with the gold standard omnibus test (candidate duplicates identified by any approach and confirmed following human scrutiny), AsySD falsely labelled 101 citations as duplicates, and human reviewers had falsely labelled 164 citations as duplicates. This gives specificity, across all 5 datasets, of 0.9991 for ASySD and 0.9986 for human reviewers; and sensitivity of 0.9775 for ASySD and 0.9522 for human reviewers.

**Figure 1:**
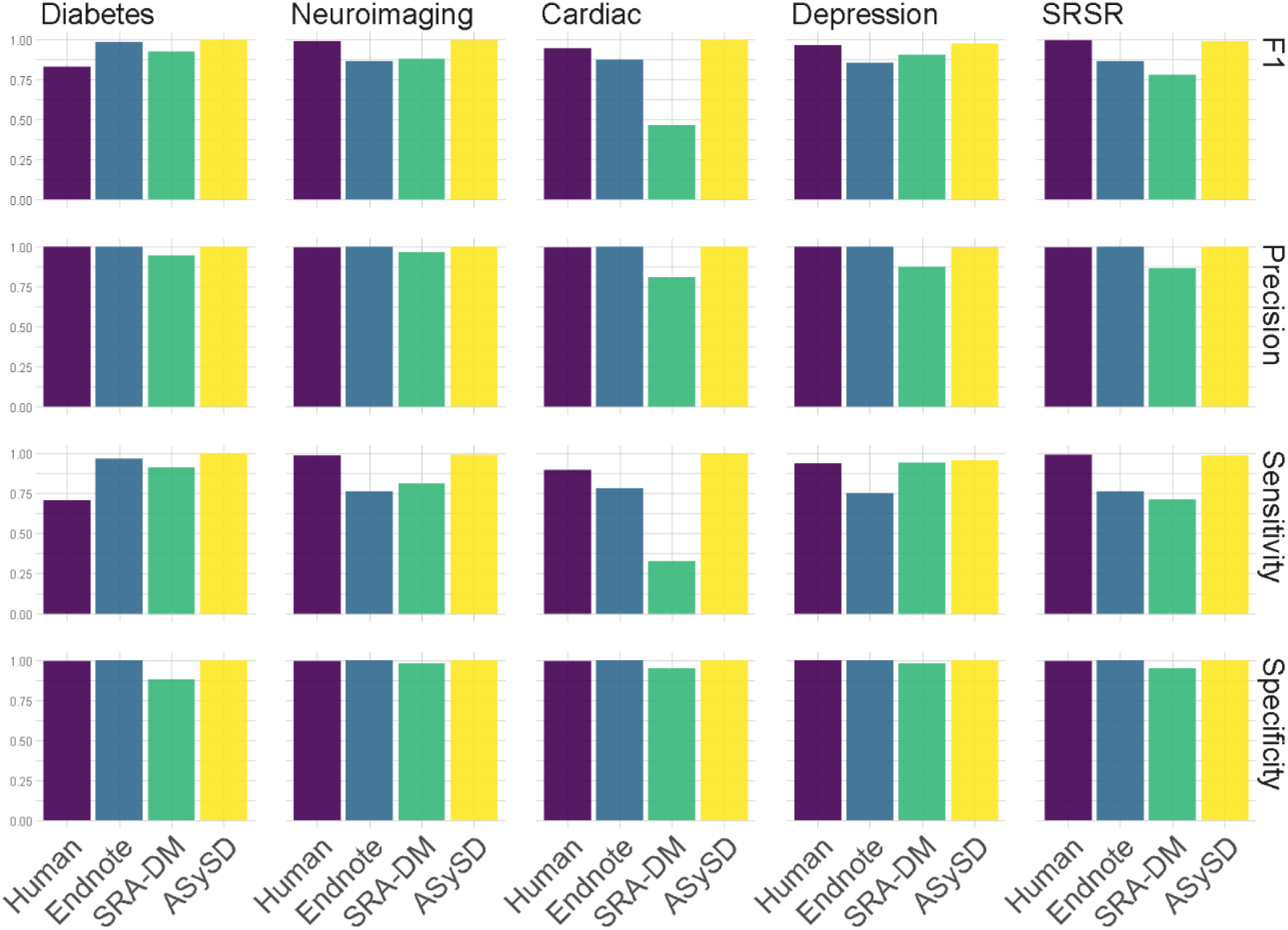
Overall performance of deduplication methods.

## Discussion

### Human error

We evaluated the performance of different deduplication approaches using datasets from past and existing systematic review projects that were not specifically established to test a deduplication tool. The rationale by which a reviewer removed any given citation is therefore not clear and there are a number of possible reasons: accidental deletion, removal due to knowledge that article was not relevant, or corrupted files. The process is also likely to be influenced by differences in how reviewers determine what a duplicate is. Often, false positives seemed to be the result of very similar publications (e.g. same title, author, and year) of which one may be a conference abstract and the other a publication. Information on what would be classed as a “duplicate” was only present in one of the corresponding gold-standard search protocols/publications. For the depression review (38), publications identified in the systematic search which reported the same primary data were considered duplicates, which diverges from our definition.

### Dataset Variability

We aimed to test each tool on heterogenous search datasets (i.e. size and number of duplicates) to determine which tool may work best for different types of systematic reviews. Endnote’s lack of sensitivity was not immediately apparent on the smallest dataset (Diabetes) but was clearly shown in larger datasets. With the exception of the Diabetes dataset, the sensitivity and specificity of Endnote was fairly consistent across all datasets. The sensitivity and specificity of ASySD was also consistent, indicating that size of dataset and duplicate proportion do not seem to affect performance. SRA-DM varied in performance, with no clear explanatory pattern emerging.

### False Positives

While ASySD and Endnote maintained low false positive rates, SRA-DM had a much larger false-positive rate. The SRA-DM was developed on clinical systematic review search datasets, which may differ in key matching criteria or other characteristics. Furthermore, it was previously assessed on 4 relatively small (by preclinical standards) datasets of fewer than 2,000 citations, which may have masked the issue. However, we did not observe trends to suggest that performance was better in smaller datasets compared to larger datasets. We noticed that citations were often removed where they were recorded as having the same DOI. This can occur when a publisher assigns a single DOI to a collection of for instance conference abstracts. In such instances, inspection of the title showed that the works were clearly independent, and not duplicates. The datasets where this was the biggest problem were also the datasets with the highest proportion of duplicates (Cardiac and Diabetes datasets).

### Time taken to remove duplicates

Endnote and ASySD were the fastest methods of deduplication, with all datasets taking under an hour to complete. SRA-DM was extremely slow for larger datasets. However, the interface was user-friendly and if a reviewer is not short of time, the program can run easily in the background without demanding too much processing power.

### Limitations OF ASySD

ASySD was developed exclusively using preclinical systematic review datasets. One dataset tested here (Neuroimaging dataset) had both clinical and preclinical studies, however performance has not been evaluated thoroughly on systematic searches within other review areas.

Due to the matching algorithm, the accuracy of ASySD is highly dependent on the quantity and quality of citation information. All systematic search datasets were from recent reviews, and although each did contain older citations, the amount of missing information was relatively low. It is unclear how any of the tools would perform on older searches or citations without page numbers, DOIs, ISBNs, and other useful bibliographic information. In these cases, it is likely that the code may need to be adapted or that the user would have to rely more heavily on manual deduplication.

Furthermore, ASySD users are likely to have different criteria for determining what counts as a “duplicate”. In future versions of ASySD, we plan to build in additional user-defined options to specify whether the algorithm should consider conference abstracts, preprints and journal articles with very similar bibliographic information to be duplicates or not. In time, with machine-readable full-text PDFs, it may also be possible to detect the same data published across multiple publications and flag these as duplicates.

While specificity was comparable to human performance, ASySD did remove some citations incorrectly. For smaller reviews in particular, this risk may not be acceptable, and future versions of ASySD will include the option to manually inspect candidate duplicates.

A key limitation of using ASySD for larger datasets (>50,000 citations) is that the processing requirements outstrip those available in our shinyapps.io subscription. We recommend that users run the application locally in R Studio for this purpose, but understand that this may cause problems for those who are not proficient in R. We are currently exploring alternative approaches which would provide sufficient processing efficiency, such as the development of deduplication software which could be installed locally. We expect that ASySD will be provided on such a platform in the near future. In the meantime, all underlying code for ASySD is openly available and has been formalised into an R package, to ensure it is interoperable and convenient for researchers wishing to integrate ASySD into their own automated evidence synthesis workflows.

## Conclusions

Across five preclinical systematic search datasets of varying size and duplicate proportions, the ASySD tool outperformed the SRA-DM and Endnote in detecting duplicates and had a false-positive rate comparable to human performance. For preclinical systematic reviews, automated duplicate removal using ASySD is a highly sensitive, reliable, and time-saving approach. The ASySD tool is freely available online via a Shiny web application and the code behind the application is open source. Further research is needed to fully evaluate and disseminate the performance of various deduplication methodologies and prioritise areas for improvement.

## Declarations

### Ethics approval and consent to participate

Not applicable

### Consent for publication

Not applicable

### Availability of data and materials

The systematic search datasets and analysis code used during the current study are available on the open science framework in the Automated Systematic Search Deduplication tool (ASySD) repository, DOI: 10.17605/OSF.IO/C9EVS

### Authors’ contributions

KH wrote the code behind the ASySD tool and devised the evaluation procedure. ZB de-duplicated the systematic review of systematic reviews dataset. KH, ES, MM, ZB and JL contributed to study concept and design.

## Acknowledgements

We would like to thank Kim Wever, Darlyn Ranis, Emily Wheater, and Alexandra Bannach-Brown for providing the gold standard de-duplicated systematic search datasets used in this study.

## References

1. Paul L, Michael R, Daniel T. The Contributions of MEDLINE, Other Bibliographic Databases and Various Search Techniques to NICE Public Health Guidance. Evidence Based Library and Information Practice. 2015;10(1).

2. Qi X, Yang M, Ren W, Jia J, Wang J, Han G, et al. Find Duplicates among the PubMed, EMBASE, and Cochrane Library Databases in Systematic Review. PLOS ONE. 2013;8(8):e71838.

3. Royle P, Milne R. Literature searching for randomized controlled trials used in Cochrane reviews: rapid versus exhaustive searches. Int J Technol Assess Health Care. 2003;19(4):591–603.

4. von Elm E, Poglia G, Walder B, Tramèr MR. Different Patterns of Duplicate PublicationAn Analysis of Articles Used in Systematic Reviews. JAMA. 2004;291(8):974–80.

5. Abraham P. Duplicate and salami publications. J Postgrad Med. 2000;46(2):67–9.

6. Huston P, Moher D. Redundancy, disaggregation, and the integrity of medical research. The Lancet. 1996;347(9007):1024–6.

7. Qi X-S, Bai M, Yang Z-P, Ren W-R. Duplicates in systematic reviews: A critical, but often neglected issue. World Journal of Meta-Analysis. 2013;1(3):97–101.

8. Tramer MR, Reynolds DJ, Moore RA, McQuay HJ. Impact of covert duplicate publication on meta-analysis: a case study. Bmj. 1997;315(7109):635–40.

9. Kwon Y, Lemieux M, McTavish J, Wathen N. Identifying and removing duplicate records from systematic review searches. Journal of the Medical Library Association : JMLA. 2015;103(4):184–8.

10. Jiang Y, Lin C, Meng W, Yu C, Cohen AM, Smalheiser NR. Rule-based deduplication of article records from bibliographic databases. Database. 2014;2014.

11. Hupe M. EndNote X9. Journal of Electronic Resources in Medical Libraries. 2019;16(3–4):117–9.

12. Bramer WM, Giustini D, de Jonge GB, Holland L, Bekhuis T. De-duplication of database search results for systematic reviews in EndNote. J Med Libr Assoc. 2016;104(3):240–3.

13. Rathbone J, Carter M, Hoffmann T, Glasziou P. Better duplicate detection for systematic reviewers: evaluation of Systematic Review Assistant-Deduplication Module. Systematic Reviews. 2015;4(1):6.

14. Westgate MJ. revtools: An R package to support article screening for evidence synthesis. Research synthesis methods. 2019;10(4):606–14.

15. Smalheiser NR, Lin C, Jia L, Jiang Y, Cohen AM, Yu C, et al. Design and implementation of Metta, a metasearch engine for biomedical literature retrieval intended for systematic reviewers. Health Inf Sci Syst. 2014;2.

16. Jiang Y, Lin C, Meng W, Yu C, Cohen AM, Smalheiser NR. Rule-based deduplication of article records from bibliographic databases. Database (Oxford). 2014;2014:bat086.

17. Mueen Ahmed K, Al Dhubaib B. Zotero: A bibliographic assistant to researcher. Journal of Pharmacology and Pharmacotherapeutics. 2011;2(4):303–5.

18. Zaugg H, West RE, Tateishi I, Randall DL. Mendeley: Creating communities of scholarly inquiry through research collaboration. TechTrends. 2011;55(1):32–6.

19. de Vries RB, Wever KE, Avey MT, Stephens ML, Sena ES, Leenaars M. The usefulness of systematic reviews of animal experiments for the design of preclinical and clinical studies. ILAR journal. 2014;55(3):427–37.

20. Sena ES, Currie GL, McCann SK, Macleod MR, Howells DW. Systematic reviews and meta-analysis of preclinical studies: why perform them and how to appraise them critically. Journal of cerebral blood flow and metabolism : official journal of the International Society of Cerebral Blood Flow and Metabolism. 2014;34(5):737–42.

21. Lorenzetti DL, Ghali WA. Reference management software for systematic reviews and meta-analyses: an exploration of usage and usability. BMC Medical Research Methodology. 2013;13(1):141.

22. Bramer WM, Giustini D, de Jonge GB, Holland L, Bekhuis T. De-duplication of database search results for systematic reviews in EndNote. Journal of the Medical Library Association : JMLA. 2016;104(3):240–3.

23. Elliott J, Turner T, Clavisi O, Thomas J, Higgins J, Mavergames C. Living systematic reviews: an emerging opportunity to narrow the evidence-practice gap. PLoS Med. 2014;11.

24. Bashir R, Surian D, Dunn AG. Time-to-update of systematic reviews relative to the availability of new evidence. Systematic Reviews. 2018;7(1):195.

25. Hair K, Bahor Z, Macleod M, Sena E. Protocol: evaluating the performance of automated deduplication tools for systematic reviews. Open Science Framework.

26. Borg MSaA. {The RecordLinkage Package: Detecting Errors in Data. The R Journal. 2010;2(2):61–7.

27. Currie GL, Angel-Scott H, Colvin L, Cramond F, Hair K, Khandoker L, et al. Animal models of chemotherapy-induced peripheral neuropathy: a machine-assisted systematic review and meta-analysis A comprehensive summary of the field to inform robust experimental design. bioRxiv. 2018:293480.

28. McCann SK. Antioxidants - focal ischaemia 2018 [Available from: https://app.syrf.org.uk/projects/153e59fe-daa2-43db-8a43-fd9e01d650e3/detail

29. Simonato M, Iyengar S, Brooks-Kayal A, Collins S, Depaulis A, Howells DW, et al. Identification and characterization of outcome measures reported in animal models of epilepsy: Protocol for a systematic review of the literature-A TASK2 report of the AES/ILAE Translational Task Force of the ILAE. Epilepsia. 2017;58 Suppl 4:68–77.

30. Hair K. ASySD 2021 [Available from: https://github.com/camaradesuk/ASySD.

31. Hair K. ASySD web application 2019 [Available from: https://camarades.shinyapps.io/RDedup/.

32. Hair K. ASySD_shiny [Web Page]. 2020 [Available from: https://github.com/camaradesuk/ASySD_shiny.

33. Wever K, Ranis D, Hooijmans C, Riksen N. The effects of the novel anti-diabetic drugs SGLT2i, GLP-1a and DPP4i on atherosclerosis - A systematic review and meta-analysis of animal studies. PROSPERO 2018 CRD42018116259. 2018.

34. Wheater ENW, Stoye DQ, Cox SR, Wardlaw JM, Drake AJ, Bastin ME, et al. DNA methylation and brain structure and function across the life course: a systematic review. Neuroscience & Biobehavioral Reviews. 2020.

35. Wever KE, Hooijmans CR, Riksen NP, Sterenborg TB, Sena ES, Ritskes-Hoitinga M, et al. Determinants of the Efficacy of Cardiac Ischemic Preconditioning: A Systematic Review and Meta-Analysis of Animal Studies. PLOS ONE. 2015;10(11):e0142021.

36. Bannach-Brown A, Liao J, Wegener G, Macleod M. Understanding in vivo modelling of depression in non-human animals: a systematic review protocol. Evidence-based Preclinical Medicine. 2016;3(2):e00024.

37. Hair K, McCann S. Protocol for a systematic review of preclinical systematic reviews. Open Science Framework. 2020.

38. Alexandra BB, Jing L, Gregers W, Malcolm M. Understanding in vivo modelling of depression in non-human animals: a systematic review protocol. Evidence-based Preclinical Medicine. 2016;3(2):e00024.

